# An Engineered Halotolerant Chimeric T7 RNA Polymerase for High-Yield, Low-Immunogenicity Synthesis of RNA via Simple Batch Transcription

**DOI:** 10.64898/2026.05.13.724829

**Authors:** Yuan Fang, Yan Sun, Congcong Wang, Xiaodie Wang, Lingling Zhang, Meiqi Shao, Jianchen Yu, Yiwen Liang, Qijun Qian, Pingjing Zhang

**Author notes:** CORRESPONDENCE Pingjing Zhang:, Qijun Qian.

## Abstract

The rapid advancement of mRNA therapeutics has imposed stringent requirements on both the quality and scalability of in vitro transcription (IVT) products. However, the accumulation of double-stranded RNA (dsRNA) byproducts and 3’-terminal heterogeneity during T7 RNA polymerase (T7 RNAP)-mediated transcription can robustly trigger deleterious innate immune responses and compromise translation efficiency. Existing enzyme engineering strategies frequently struggle to reconcile the trade-offs between salt tolerance, volumetric productivity, and product purity. Here, we report a novel engineering strategy for halotolerant T7 RNAP by fusing optimized mutant polymerases with diverse DNA-binding domains (e.g., Sso7d, MC1). This approach orchestrated the development of a series of chimeric T7 RNAP mutants designed to bolster catalytic activity and template selectivity under high-salt conditions while concurrently suppressing RNA-dependent RNA polymerase (RdRP) activity. Our lead chimeric mutants exhibited exceptional salt tolerance and processivity in the presence of up to 270 mM NaCl. Notably, these mutants significantly diminished dsRNA formation to less than 0.001%, while markedly improving transcript integrity and 3’ homogeneity, thereby facilitating superior translation efficiency for both linear mRNA and circular RNA (circRNA). Crucially, this heightened salt tolerance does not necessitate a trade-off in RNA yield, affording broader flexibility for downstream process optimization. In an enzymatic circRNA synthesis system, these mutants enabled a non-fed-batch configuration with high initial rNTP concentrations (15 mM each), resulting in a 50% increase in yield and achieving an unprecedented titer of 15 mg/mL. This research provides a robust enzymological solution that harmonizes quality and productivity for the industrial-scale manufacturing of high-concentration, low-immunogenicity RNA.

## Introduction

Following the success of COVID-19 vaccines, mRNA technology is rapidly diversifying from infectious disease prophylaxis toward personalized oncology vaccines and protein replacement therapies. The inherent sequence programmability and modularity of mRNA platforms afford exceptional therapeutic flexibility^1–5^. However, the clinical transition toward high-dose regimens and chronic treatment modalities poses formidable challenges to product quality and scalable manufacturing capacity^6–10^. The Critical Quality Attributes (CQAs) of mRNA— encompassing the accumulation of double-stranded RNA (dsRNA) byproducts, 3’-terminal heterogeneity, and overall transcript integrity—are paramount^11,12^. In particular, dsRNA contaminants generated via antisense transcription or self-primed extension during in vitro transcription (IVT) function as potent agonists for pattern recognition receptors (PRRs) such as MDA5 and RIG-I. These impurities can robustly trigger hyperactivation of the innate immune response, culminating in dose-limiting toxicities and the profound suppression of translation efficiency^13–17^.

Current optimization of mRNA synthesis revolves around two primary trajectories: the refinement of IVT reaction parameters and the engineering of RNA polymerase functionalities^9,18–22^. Notwithstanding substantial advancements, the engineering of T7 RNA polymerase (T7 RNAP) remains constrained by several technical bottlenecks^18–20^. To date, protein engineering strategies aimed at mitigating double-stranded RNA (dsRNA) accumulation during T7 RNAP-mediated transcription have generally followed three distinct paradigms. The first paradigm utilizes site-directed mutagenesis to attenuate promoter-independent initiation or facilitate promoter clearance, thereby curtailing dsRNA synthesis^19,20,23,24^. However, such mutants frequently exhibit pronounced salt sensitivity, which constrains the optimization of rNTP concentrations and buffer ionic strength, ultimately limiting volumetric productivity^20^. The second approach employs directed evolution to yield thermostable mutants capable of operating at elevated temperatures (45–50°C), thereby destabilizing RNA secondary structures and reducing dsRNA formation^25–29^. Nevertheless, achieving the desired dsRNA threshold often necessitates extreme thermal conditions, which may inadvertently compromise transcript integrity through heat-induced RNA degradation^25,26^. The third strategy focuses on domain fusion (e.g., Sso7d) or the development of biotin-streptavidin-based covalent anchoring systems to stabilize the enzyme-template complex under high-salt conditions^30–33^. These modifications aim to suppress RNA-dependent RNA polymerase (RdRP) activity by robustly enhancing affinity for the DNA template. While domain-fused RNA polymerases have been shown to partially inhibit antisense RNA synthesis in high-salt environments, the underlying molecular mechanisms remain to be fully elucidated. Such insights are critical for designing mutants with sufficient salt resilience to maintain high titers while simultaneously minimizing dsRNA byproducts. Furthermore, co-tethering systems necessitate complex, multi-step procedures, which may not represent the optimal solution for industrial-scale upscaling and standardized manufacturing. Consequently, the development of a T7 RNAP mutant that exhibits superior salt tolerance and heightened catalytic activity while maintaining high volumetric productivity remains an urgent desideratum in the field of RNA therapeutics.

Here, we propose a bifurcated engineering framework distinct from previously reported paradigms, wherein a repertoire of chimeric T7 RNAP mutants was constructed by integrating combinatorial mutations with heterologous DNA-binding domains (DBDs) fused to the N-terminal domain (NTD) (Figure 1) This strategy was designed to synergistically bolster catalytic activity under high-salt conditions, ensuring the robust suppression of dsRNA byproducts while simultaneously maximizing volumetric RNA productivity. Our lead chimeric mutants exhibited exceptional performance metrics, characterized by a significant reduction in dsRNA accumulation (<0.001%), enhanced transcript integrity, and improved 3’ homogeneity. Importantly, this robust salt resilience does not necessitate a trade-off in RNA yield, thereby affording broader operational flexibility for downstream process optimization. Within an enzymatic circular RNA (circRNA) synthesis platform, these mutants facilitated a non-fed-batch configuration utilizing elevated initial rNTP concentrations (15 mM each), resulting in a 50% increase in yield and achieving an unprecedented titer of 15 mg/mL. Collectively, this research elucidates a potent enzymological solution that harmonizes product quality and yield, offering a scalable solution for the industrialized manufacturing of high-concentration, low-immunogenicity RNA therapeutics.

**Figure 1.**
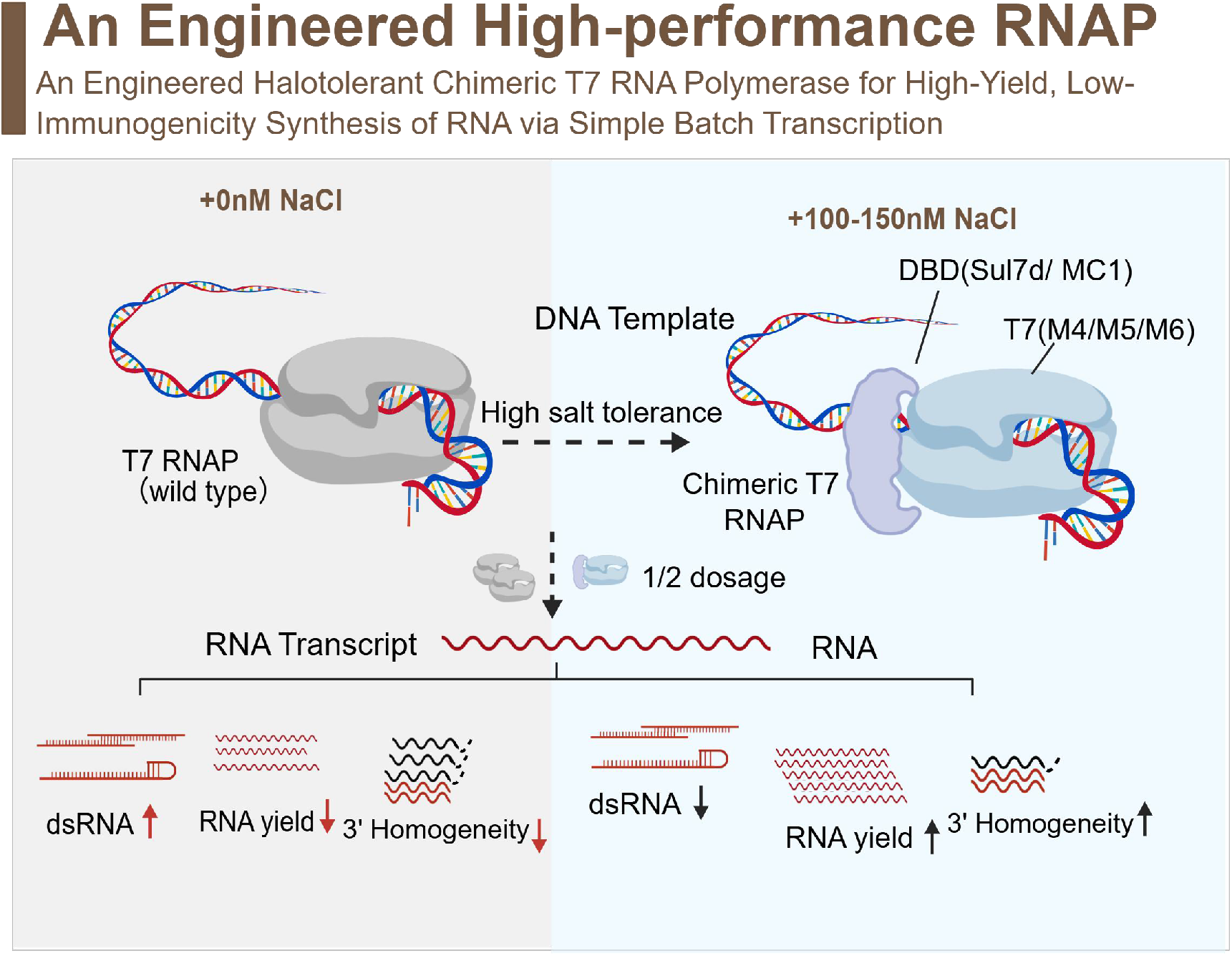
graphical abstract. Schematic comparing wild-type and chimeric T7 RNAP for high-salt IVT. The DNA-binding domain (DBD)-fused mutants (T7 M4/M5/M6; Sso7d/MC1), utilized at a 50% reduced dosage, demonstrated superior RNA titers compared to the wild-type enzyme. Under 100–150 mM NaCl, these chimeric polymerases significantly mitigated dsRNA byproduct formation and enhanced 3’ homogeneity. This optimized catalytic profile facilitates high-efficiency, “one-pot” synthesis of both conventional mRNA and circRNA.

## Results

### Rational Design of Halotolerant T7 RNA Polymerase Chimeras via Catalytic Core and DBD Integration

Bacteriophage 7 RNAP exhibits profound sensitivity to high-salt environments, wherein total salt concentrations exceeding 150 mM typically trigger severe transcriptional inhibition. Extensive prior investigations have demonstrated that the introduction of specific amino acid substitutions can significantly bolster the thermostability and thermal activity of the enzyme^34^. Given the intrinsic linkage between T7 RNAP stability and environmental resilience, we hypothesized that mutations capable of stabilizing the spatial conformation of the enzyme might concurrently confer heightened salt tolerance. Accordingly, by leveraging and integrating several canonical thermostabilization strategies, we engineered a repertoire of T7 RNAP mutants, designated as M4, M5, and M6 (Figure 2A, Supplementary File 2: Table S1).

**Figure 2.**
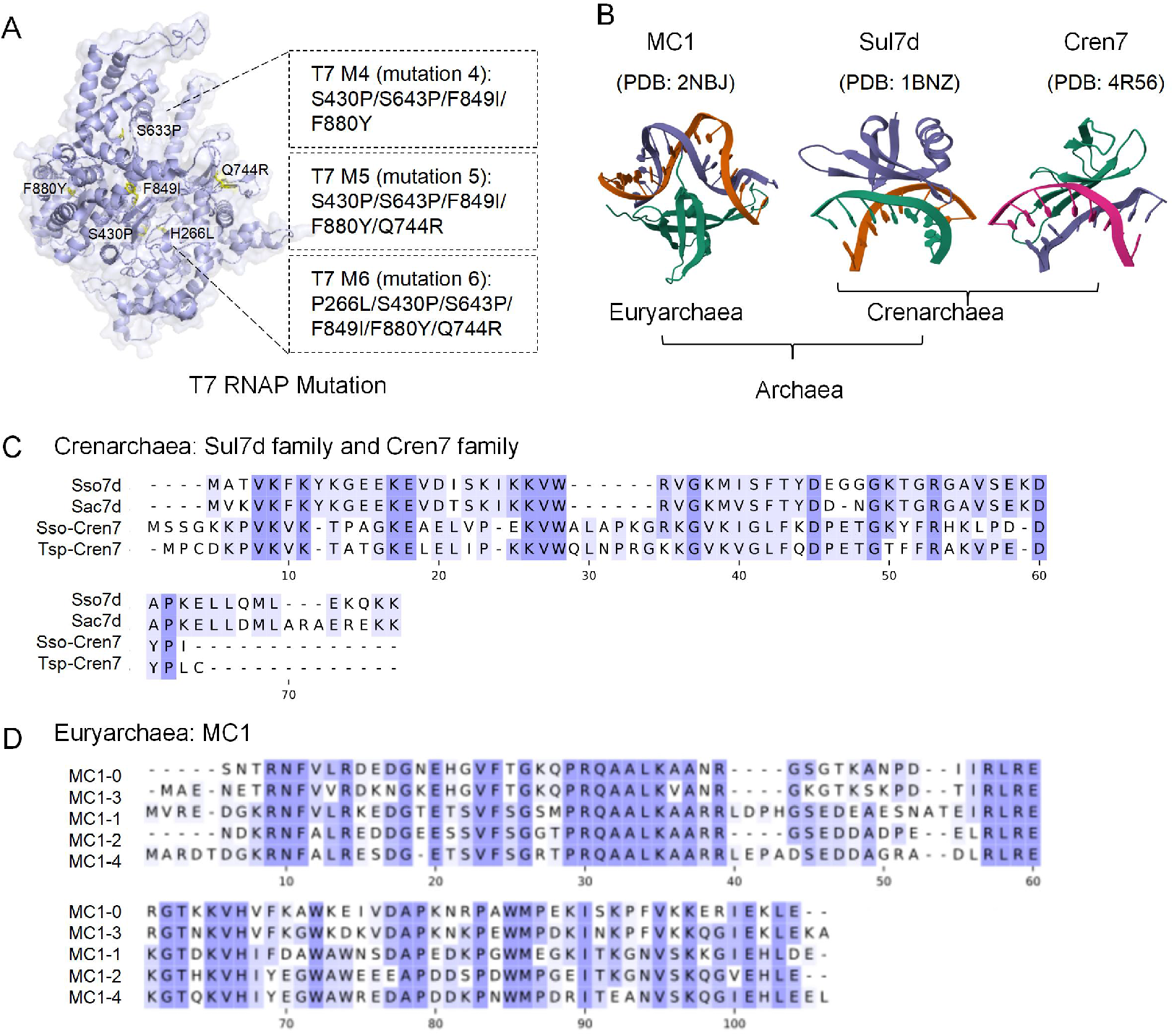
Structural architecture and phylogenetic analysis of engineered T7 RNAP mutants and archaeal DBDs. (A) Structural mapping of amino acid substitutions within the T7 RNAP mutants M4-M6. The M4 scaffold comprises the quadruple mutation S430P/S643P/F849I/F880Y; the M5 mutant incorporates an additional Q744R substitution; and the M6 iteration further includes the P266L mutation. (B) Representative three-dimensional conformations of candidate archaeal DBDs derived from Euryarchaeota (MC1, PDB: 2NBJ; Sul7d, PDB: 1BNZ) and Crenarchaeota (Cren7, PDB: 4R56). (C, D) Multiple sequence alignments of (C) Sul7d- and Cren7-family orthologs from Crenarchaeota and (D) Euryarchaeota-derived MC1 mutants. Conserved residues are highlighted to indicate evolutionary conservation and potential structural significance.

Recent investigations have elucidated that chimeric T7 RNAP harboring Sso7d—a small archaeal DNA-binding protein of the Sul7d (Sulfolobus 7kDa DNA-binding protein) family— robustly enhances salt tolerance by strengthening interactions with the double-stranded DNA (dsDNA) template and its promoter^30^. Furthermore, this fusion modulates the enzyme’s RdRP activity, thereby facilitating a reduction in dsRNA byproducts. We postulate that the intrinsic binding affinity of diverse DBDs for the DNA template represents a primary determinant of the chimeric polymerase’s resilience to high-salt environments, ultimately dictating both RNA yield and transcript quality. Archaeal genomes encode several classes of architectural DNA-bending proteins^35,36^, including the Sul7d, Cren7(Crenarchaeal 7 kDa)^37^, and MC1(Methanogen Chromosomal protein 1) ^38^ families (Figure 2B). Structural characterizations indicate that Sul7d and Cren7 associate with the convex surface of bent DNA, whereas members of the MC1 family interact with the concave surface. In archaea, MC1, Sul7d, and Cren7 function as pivotal, non-sequence-specific chromatin-organizing monomers^37^. Specifically, Sul7d and Cren7, predominantly found in the order Sulfolobales, utilize their surface β-sheets to anchor within the DNA minor groove^39^. Conversely, MC1 is primarily identified in methanogenic archaea; notably, its interaction with DNA has been reported to be transiently attenuated as ionic strength increases^39^. Building upon these insights, we screened four Sul7d/Cren7-derived peptides (Figure 2C, Supplementary File 2: Table S1) and five MC1-derived peptides (Figure 2D, Supplementary File 2: Table S1) for fusion with the NTD of T7 RNAP or its mutants.

### N-terminal Fusion of Heterotypic DNA-Binding Domains Empowers T7 RNA Polymerase with Enhanced Specific Activity and High-Salt Robustness

Utilizing the fluorogenic interaction between DFHBI and the iSpinach RNA aptamer, we established a real-time quantitative assay for monitoring IVT products^40–43^. Transcriptional efficiencies of various mutants were systematically evaluated by monitoring fluorescence kinetic profiles under both baseline (0 mM NaCl) and high-salt (100 mM NaCl) conditions. Using specific activity—defined as the quantity of RNA synthesized per unit of enzyme per unit time—as the primary metric, we identified candidates with optimal catalytic performance. To screen for high-efficiency T7 RNAP scaffolds suitable for Sso7d fusion, a repertoire of mutants harboring an N-terminal Sso7d domain was designed and engineered (Figure 3A). Activity assays revealed that N-terminal Sso7d fusion robustly enhanced T7 RNAP activity under high-salt conditions. Notably, the M5 mutant exhibited superior performance compared to the M4 and M6 candidates (Supplementary File 1: Figure S1A). Consequently, the T7 M5 mutant was selected as the primary scaffold for the subsequent development of chimeric mutants.

**Figure 3.**
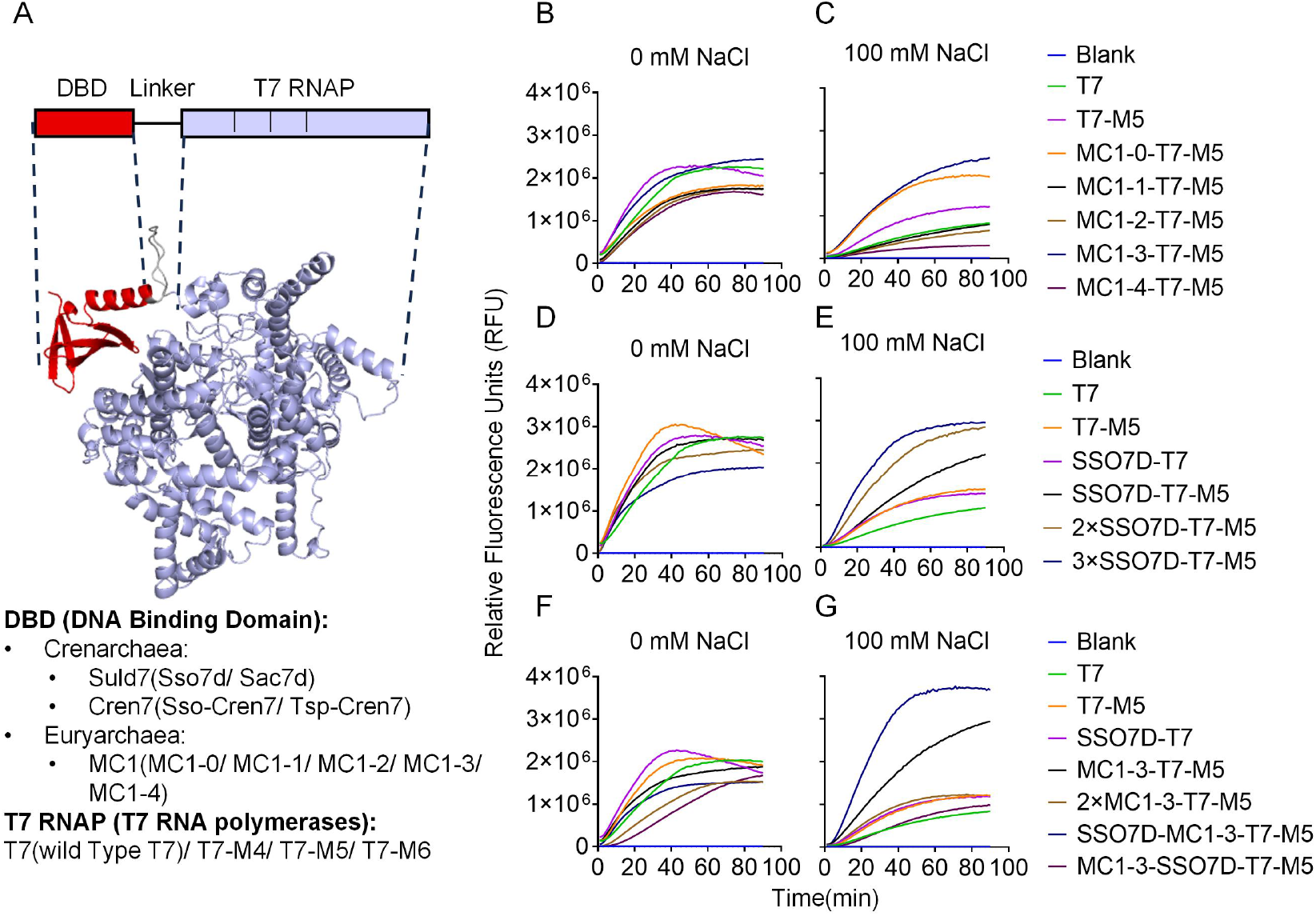
Real-time kinetic profiling of chimeric T7 RNAPs engineered with archaeal DNA-binding domains. (A) Schematic representation and structural model of the chimeric architecture, featuring an N-terminal archaeal DBD tethered to the T7 RNAP M5 scaffold via a flexible linker. Evaluated DBDs include the Sul7d/Cren7 families from Crenarchaeota and the MC1 family from Euryarchaeota. (B-G) In vitro transcription kinetics monitored via real-time fluorescence under low-salt (0 mM NaCl: B, D, F) and high-salt (100 mM NaCl: C, E, G) conditions.(B, C) Catalytic activity of MC1-mutant fusions.(D, E) Comparative performance of Sso7d-T7 and Sso7d-T7-M5 mutants across varying DBD valencies (1×, 2×, or 3× copy numbers).(F, G) Combinatorial fusions integrating MC1-3 and Sso7d. Notably, DBD-fused M5 mutants exhibit significantly enhanced halotolerance and robust transcriptional activity at 100 mM NaCl, contrasting the salt-induced inhibition observed in wild-type T7 RNAP.

To identify Sul7d-family sequences capable of enhancing enzymatic potency, we engineered a repertoire of mutants by fusing Sso7d to the NTD of both T7(wild-type) and T7-M5 RNA polymerases (Supplementary File 1: Figure S1B). Our data reveal that the T7 M5 mutation and chimeric fusions with Sul7d-family proteins (Sso7d and Sac7d) robustly augmented transcriptional activity. Notably, these two strategies orchestrated a synergistic effect, wherein Sso7d- or Sac7d-chimeric T7 M5 mutants achieved peak transcriptional performance— approximately twofold greater than that of the T7 RNAP. In stark contrast to the non-chimeric T7 RNAP, the two Cren7-fused polymerases failed to exhibit any discernible activity enhancement and, instead, displayed significantly impaired transcriptional capacity (Supplementary File 1: Figure S1B).

Similarly, a repertoire of mutants was engineered to evaluate the impact of MC1-family peptides on T7 RNAP catalytic activity (Figure 3A). Our data reveal that none of the MC1-domain fusions with the T7 RNAP M5 mutant yielded further enhancements in enzymatic activity under low-salt conditions; notably, fusions with MC1-0, MC1-1, MC1-2, and MC1-4 even exhibited a marginal decline in performance (Figure 3B). Under high-salt conditions, the T7 RNAP M5 mutant displayed an approximately twofold increase in transcriptional activity compared to the T7 RNAP. Furthermore, the incorporation of DBDs derived from MC1-0 and MC1-3 augmented the specific activity of the T7 RNAP M5 scaffold by an additional twofold. Remarkably, the fluorescence intensity generated by their RNA aptamer products was comparable to that of T7 RNAP under low-salt conditions (Figure 3C), with the MC1-3 chimeric mutant emerging as the top performer. These disparate results suggest that T7 RNAP harboring N-terminal MC1 fusions may inherit the adaptive responses of their parental archaea to varying ionic strengths. Upon elevation of environmental salt concentrations, the robustly positively charged surface of the MC1 domain maintains electrostatic attraction to the DNA phosphate backbone. This interaction effectively compensates for the attenuated affinity between the T7 RNAP and the DNA template, thereby facilitating higher processivity and maintaining the stability of the DNA-RNA hybrid under high-salt conditions.

Given the divergent DNA-binding interfaces and architectural bending mechanisms inherent to the Sul7d and MC1 families, we engineered several chimeric T7 RNAP mutants featuring tandem DBDs. These architectures encompassed homozygous tandem repeats (2–3 copies of proteins from the same family) as well as heterologous combinations of MC1 and Sul7d family members, with all proteins successfully purified for subsequent characterization (Supplementary File 1: Figure S2). Under low-salt conditions, neither the homozygous nor the heterologous tandem strategies significantly augmented the specific activity of the non-chimeric T7 M5. Conversely, a majority of these chimeric mutants displayed diminished transcriptional activity, with the sole exception of a marginal enhancement observed for Sso7d-T7 M5 (Figures 3D & 3F). In contrast, under high-salt conditions, the M5 mutants harboring 2–3 copies of Sso7d robustly achieved peak transcriptional potency, exhibiting a clear copy-number-dependent enhancement in transcriptional activity (Figure 3E). Intriguingly, for the MC1-3 family, doubling the DBD copy number resulted in a decline in high-salt activity compared to the single-copy chimeric mutant (Figure 3G). The heterologous tandem fusions yielded disparate performance profiles; notably, Sso7d-M1-3-T7 M5 displayed the highest activity, whereas the reversed configuration, M1-3-Sso7d-T7 M5, performed comparably to the baseline T7 M5 (Figure 3G). These divergent outcomes likely elucidate the complex synergistic interplay between linker-mediated flexibility and the distinct DNA-bending geometries of the fused domains. Such variations in tandem organization may differentially modulate the stability of the DNA-RNA hybrid within the T7 RNAP, thereby dictating the overall catalytic processivity and salt resilience of the chimeric enzymes.

In summary, the N-terminal integration of Sul7d and MC1-derived peptides establishes a robust framework for markedly augmenting both the specific activity and halotolerance of T7 RNAP. Our findings underscore that chimeric mutants featuring tandem Sso7d repeats or the Sso7d-MC1-3 heterologous configuration exhibit superior catalytic potency and structural stability under high-salt conditions (100 mM NaCl).

### Tandem DNA-Binding Domains and High-salt Synergistically Improve RNA 3’ Homogeneity

During the IVT of extended mRNA transcripts, 3’ heterogeneity represents a ubiquitous phenomenon, primarily arising from the non-templated terminal extension activity inherent to T7 RNAP. This heterogeneity establishes the structural scaffold for the formation of dsRNA. Upon termination, 3’ termini harboring adventitious nucleotides may undergo intramolecular base-pairing to form hairpin structures or transiently hybridize with extrinsic RNA strands or template fragments via intermolecular interactions. Subsequently, T7 RNAP leverages its intrinsic RdRP activity to utilize these RNA-RNA hybrids as substrates for further extension, ultimately generating hairpin-like or linear dsRNA byproducts (Figure 4A). To elucidate whether our engineered high-activity T7 RNAP mutants achieve superior 3’ homogeneity, we synthesized sense and antisense long single-stranded DNA (ssDNA) oligonucleotides to reconstruct the transcription templates described by Dousis et al. ^20^ RNA transcripts were prepared and subsequently subjected to an RNase T1-based analytical workflow, as previously established by Jiang et al. ^44^, to quantitatively assess 3’ homogeneity.

**Figure 4.**
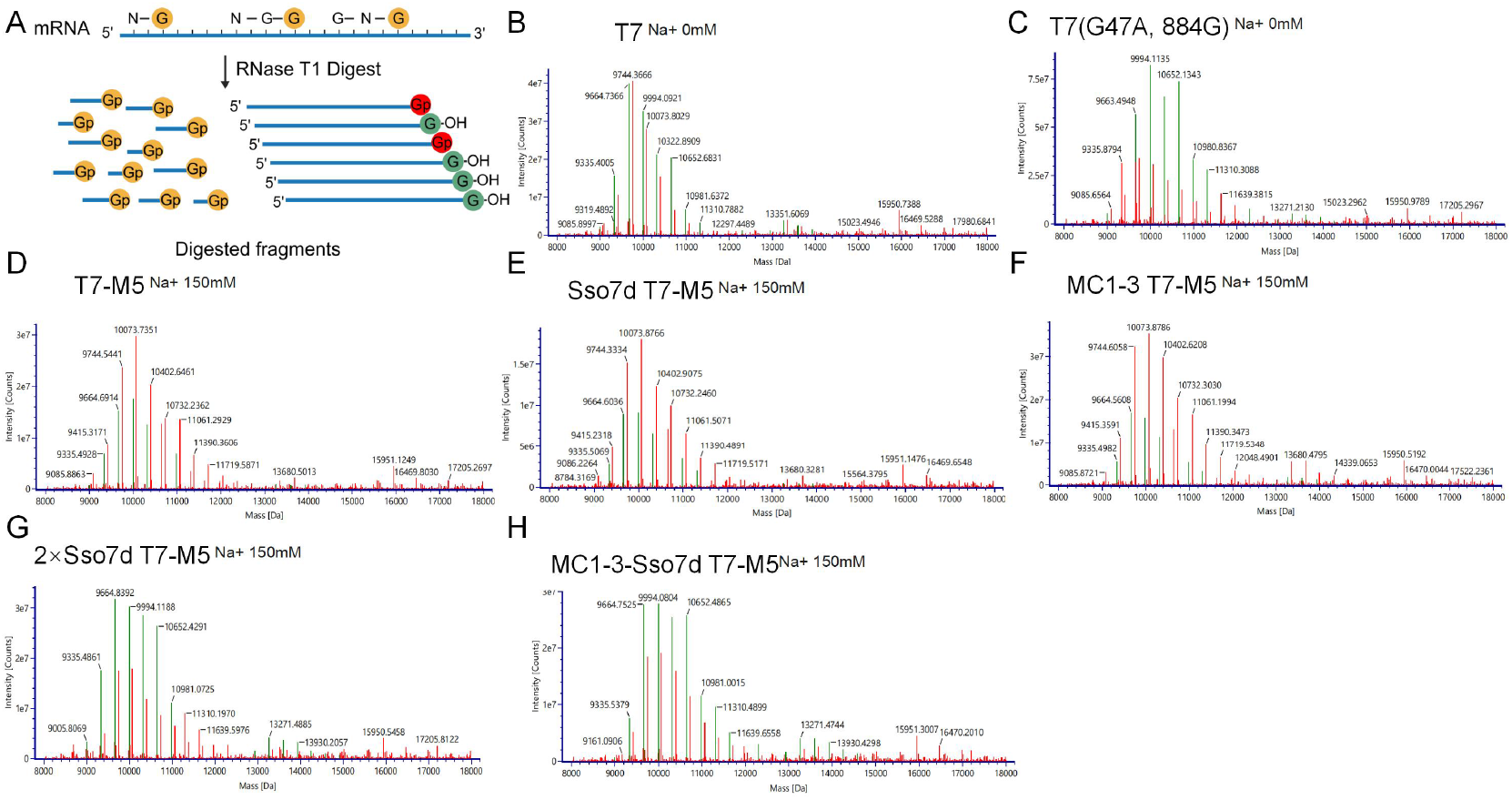
Mass spectrometric characterization of IVT products demonstrating that DBD-fusions improve 3’ homogeneity under high-salt. (A) Schematic representation of RNase T1 Digest Assay. (B, C) Mass spectrometric profiles of RNA synthesized by T7 RNAP (B) and the T7(G47A/P884G) mutant (C) under low-salt conditions (0 mM NaCl).(D–H) Mass spectra acquired at 150 mM NaCl for the T7-M5 scaffold (D) and the engineered chimeras: Sso7d-T7-M5 (E), MC1-3-T7-M5 (F), 2×Sso7d-T7-M5 (G), and MC1-3-Sso7d-T7-M5 (H).Relative to both the WT enzyme and the M5 mutant, the chimeric fusions (E–H) exhibit significantly enhanced transcriptional precision, characterized by the recovery of full-length transcript integrity and the suppression of 3’ end heterogeneity despite elevated salt concentrations.

Experimental results reveal that under low-salt conditions, chimeric RNA polymerase mutants—owing to their enhanced T7 promoter binding affinity and elevated specific activity— exhibit an increased propensity for non-specific extension by associating with nascent RNA strands or DNA oligonucleotides. This phenomenon is underscored by mass spectrometry data, which characterize a distinct bimodal peak profile (Supplementary File 1: Figure S3). In contrast, high-salt transcription environments augment the ionic shielding effect, thereby attenuating the electrostatic interactions between the polymerase and the non-specific nucleic acid backbone. This transition compels the enzyme to rely exclusively on stable initiation at the promoter or the formation of an authentic elongation complex, effectively abolishing non-specific transcriptional byproducts (Figure 4). Furthermore, the high-salt environment interferes with the uptake of non-templated nucleotides within the T7 RNAP. This increased ionic strength facilitates prompt enzyme dissociation upon reaching the template terminus during termination, preventing the RNAP from lingering at the 3’ end to catalyze stochastic terminal tailing.

Interestingly, DBD fusions (Sso7d-T7-M5 and MC1-3-T7-M5) exhibited a marginal increase, rather than a decrease, in non-templated terminal extension under high-salt conditions (Figures 4E–F). In contrast, the homotypic and heterotypic tandem DBD chimeras (2×Sso7d-T7-M5 and MC1-3-Sso7d-T7-M5) markedly enhanced the binding affinity for dsDNA (Figures 4G–H). This augmented interaction presumably improves 3’ homogeneity by either modulating the elongation rate or increasing the stability of the termination complex. These results are comparable to the ≥70% 3’ homogeneity reported for the G47A/884G mutants^20^, which similarly leverage attenuated processivity to refine product uniformity. Collectively, our data underscore that chimeric RNAP mutants, when operated under high-salt conditions, markedly enhance 3’ homogeneity, thereby mitigating the immunogenic potential of synthetic mRNA.

### Synergistic Effects of Ionic Strength and chimeric T7 RNAP on mRNA Yield, Transcript Integrity, and dsRNA Suppression

To systematically assess the impact of salt concentration on the catalytic performance of chimeric T7 RNAP variants, the ionic strength contribution from rNTP sodium salts was rigorously accounted for. Calculations indicated that each 10 mM rNTP (as sodium salts) contributes approximately 120 mM background Na^+^ to the reaction milieu. By supplementing the reaction with 0, 100, and 150 mM exogenous NaCl, we established three distinct experimental conditions with cumulative ionic strengths of approximately 120, 220, and 270 mM, respectively. Concentrations of Tris-HCl and MgCl_2_ were standardized to isolate the effects of monovalent cations. Using PiggyBac (PB) transposase mRNA as a reporter, we assessed transcriptional efficiency and product quality across these gradients. Notably, the transcript purity and co-transcriptional capping efficiency remained robustly sustained even under elevated salinity regimes. The purity of the IVT products was scrutinized via chromatographic analyses (Supplementary File 1: Figure S4) and the capping efficiency was analyzed (Supplementary File 1: Figure S5).

Experimental results demonstrated that the control T7 (G47A, 884G) mutant was incapable of initiating transcription at 10 mM of each rNTP in the absence of exogenous NaCl; consequently, rNTP concentrations were reduced to 4 mM to facilitate PB mRNA synthesis (Figure 5A). Wild-type (WT) T7 RNAP exhibited acute salt sensitivity, with volumetric yields collapsing from 7.5 mg/mL to below 2 mg/mL upon the addition of 100 mM NaCl (Figure 5A-B), and declining precipitously to negligible levels at 150 mM NaCl (Figure 5C). While Sso7d-T7 RNAP sustained its activity and yield at +100 mM NaCl, it failed to withstand the +150 mM NaCl condition (Figures 5B and 5C). In contrast, the halotolerant T7 M5 scaffold and its chimeric derivatives demonstrated robust performance. Yields for T7 M5 remained stable at 150 mM NaCl, and with the exception of the MC1-3-T7 M5 mutant—which showed a minor attenuation in productivity—all chimeric T7-M5 mutants maintained high volumetric RNA yields at 270 mM cumulative ionic strength (Figure 5C). Notably, all chimeric RNAP mutants exhibited a progressive attenuation in volumetric productivity as supplemental NaCl exceeded the 150 mM threshold (corresponding to a cumulative ionic strength > 270 mM). This downward trajectory delineates the functional upper limit for sustained catalytic efficiency within these halotolerant systems, likely reflecting a salt-induced destabilization of the elongation complex or interference with nucleotide entry at extreme ionic strengths (Supplementary File 1: Figure S6). Transcript integrity, characterized by full-length product (FLP) percentages, generally improved under high-salt conditions. Our data reveal that the T7-M5 and its monomeric DBD chimeras maintained FLP values exceeding 90% at 150 mM NaCl (Figure 5G-I). The homotypic tandem 2×Sso7d-T7-M5 mutant demonstrated a slight attenuation in FLP, though it remained consistently robust at approximately 88% across all experimental conditions. Interestingly, the heterotypic tandem MC1-3-Sso7d-T7-M5 mutant exhibited significantly reduced FLP in low-salt environments but recovered to ∼88% integrity at 150 mM NaCl (Figure 5G, I). Comparatively, the control T7 (G47A, 884G) mutant exhibited a marked decline in FLP values under elevated Mg^2+^ concentrations (Figure 5G). These findings suggest that stronger DBD-dsDNA adsorption requires higher salt concentrations to achieve optimal balance, thereby attaining maximal transcriptional activity and FLP values.

**Figure 5.**
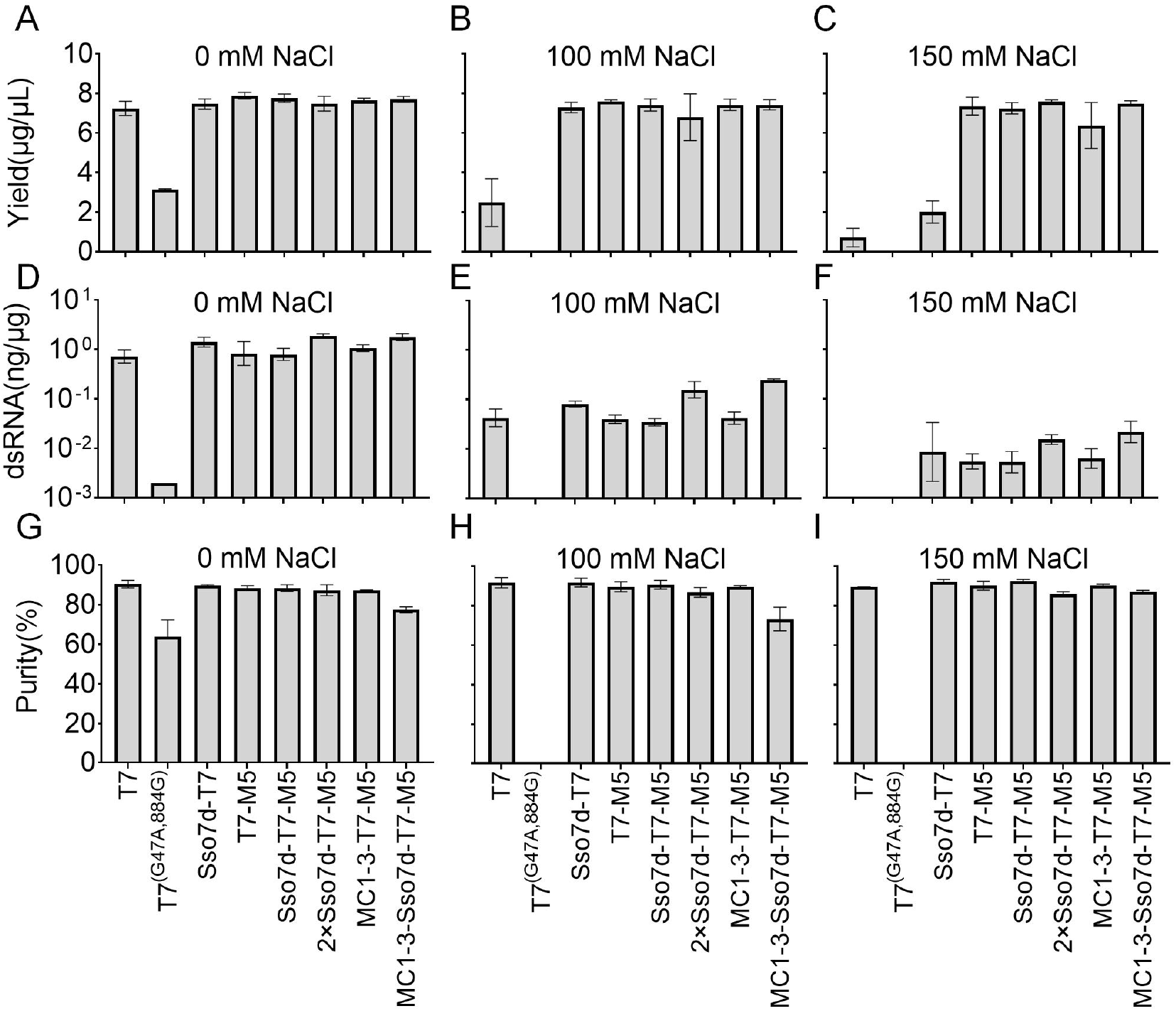
Quantitative assessment of transcriptional performance and product integrity of T7 RNAP mutants across a NaCl gradient. (A–C) Volumetric RNA yield (μg/μL), (D–F) residual dsRNA content (ng/μg total RNA, logarithmic scale), and (G–I) fractional transcript purity (%) were evaluated at 0, 100, and 150 mM NaCl. While T7 and the T7(G47A/P884G) mutant exhibited a precipitous decline in biosynthetic output under elevated ionic strength, the DBD-tethered chimeras (Sso7d-T7-M5, 2×Sso7d-T7-M5, MC1-3-T7-M5, and MC1-3-Sso7d-T7-M5) sustained high-level productivity. Notably, these chimeric mutants maintained minimal dsRNA accumulation and superior transcript homogeneity across all tested salt concentrations, effectively overcoming the salt-induced inhibition typical of conventional IVT systems. Data are expressed as mean ± s.d. (n=3).

Furthermore, our results demonstrate that all RNA polymerases exhibit a concentration-dependent reduction in dsRNA contaminants as salt levels increase within the 100-150 mM NaCl range (Figure 5D-F). At baseline (∼120 mM total Na^+^), dsRNA levels exceeded 0.1% (1 ng/μg) for all mutants (Figure 5D). Supplementation with 100 mM NaCl reduced dsRNA content in monomeric DBD chimeras (Sso7d-T7, Sso7d-T7 M5 and MC1-3-T7 M5) and T7-M5 to 0.005-0.01% (Figure 5E), further plummeting to below 0.001% at 150 mM NaCl (Figure 5F). By contrast, MC1-3-Sso7d-T7 M5 RNAP exhibited elevated dsRNA background across all tested regimes: >0.15% (1.5 ng/μg) at 0 mM NaCl, decreasing to ∼0.02% (0.2 ng/μg) and 0.002% (0.02 ng/μg) at 100 mM and 150 mM NaCl, respectively, consistently remaining higher than monomeric DBD-chimeric counterparts under equivalent conditions (Figures 5D-F). Notably, under low-salt conditions, wild-type (WT) T7 RNAP produced slightly lower dsRNA byproducts than the monomeric DBD-chimeric variants (Figure 5D). However, at 100 mM NaCl, the WT group exhibited a marginal increase in dsRNA (Figure 5B), a phenomenon likely attributable to the concentration effect following reduced unit yields. These findings underscore that the non-specific binding affinity between the DBD and the dsDNA template positively correlates with dsRNA formation. High ionic strength appears to alleviate this high-adsorption state, thereby facilitating high transcriptional efficiency while robustly maintaining minimal dsRNA byproducts. This is corroborated by gel electrophoresis, which showed that multimeric RNA species formed readily in low-salt conditions but decreased precipitously as salt concentrations increased (Supplementary File 1: Figure S6). Since intermolecular hybridization is a structural prerequisite for RdRP-mediated extension, high-salt environments mechanistically block dsRNA formation pathways by suppressing non-specific RNA-RNA and RNA-DNA interactions. In conclusion, T7 M5 and its chimeric mutants (including Sso7d-T7 M5, MC1-3-T7 M5, and 2×Sso7d-T7 M5) achieve an optimal balance between high yield, transcript integrity, and low dsRNA contamination, fulfilling the rigorous technical requirements for high-quality RNA synthesis.

### Halotolerant T7 RNAP Chimeras Produce Low-Immunogenicity mRNA with Enhanced Cellular Translational Potency

Having established that high ionic strength significantly mitigates dsRNA contamination during IVT, we proceeded to evaluate the translational potency and immunogenicity of the resulting mRNA in a cellular context (Figure 6A). Using co-transcriptionally capped eGFP mRNA (∼1 kb) and enzymatically capped PiggyBac (PB) transposase mRNA (∼2.1 kb) as reporter transcripts, we systematically compared the expression efficiency of monomeric and tandem Sso7d-chimeric mutants synthesized under varying ionic strengths (0 mM vs. 100–150 mM NaCl). Transfection of enzymatically capped PB mRNA revealed that transcripts synthesized under high-salt conditions (150 mM NaCl) consistently yielded superior protein expression across all T7 RNAP mutants. However, no statistically significant differences in transposase activity were observed among the low-salt synthesis groups. While high-salt mutants demonstrated a marginal improvement over the G47A, I884V benchmark previously reported by Dousis et al.^20^, the variance did not reach statistical significance (p > 0.05) (Supplementary File 1: Figure S7). This limited resolution likely stems from the non-linear dose-response relationship inherent to the transposase reporter system; because the system relies on the indirect activation of tdTomato fluorescence following genomic cleavage and integration, it may mask subtle differences in mRNA quality that do not cross a specific functional threshold. Evaluations of co-transcriptionally capped eGFP mRNA underscored a congruent trend, wherein high-salt (100 mM) transcripts generally outperformed those generated under low-salt conditions. Notably, the performance enhancement observed for the Sso7d-T7 M5 chimeric variant under the high-salt regimen reached statistical significance (Figure 6B), distinguishing it as a superior candidate for robust protein expression. To elucidate the correlation between residual dsRNA and mRNA-triggered innate immune activation, we quantified dsRNA mass fractions and the subsequent induction of IFN-β in THP-1 cells. Our results reveal that high-salt conditions robustly suppressed dsRNA formation across all experimental cohorts compared to baseline salinity. Notably, Sso7d-T7-M5 and T7-M5 demonstrated the most robust performance, achieving a 10-fold reduction in dsRNA content to approximately 0.03% (Figure 6D), significantly outperforming Sso7d-T7 and the tandem 2×Sso7d-T7-M5 mutant, which both remained near 0.1%. Congruently, the IFN-β expression profiles mirrored the dsRNA concentrations, showing a substantial decline in immunogenicity (Figure 6C). Transcripts produced by Sso7d-T7-M5 and T7-M5 reduced IFN-β induction to levels below the limit of detection. In contrast, other chimeric mutants maintained detectable, albeit attenuated, inflammatory signatures. These results confirm that the high-halotolerant engineering strategy effectively reconciles high-titer RNA production with the stringent purity required to evade cellular pattern recognition receptors.

**Figure 6.**
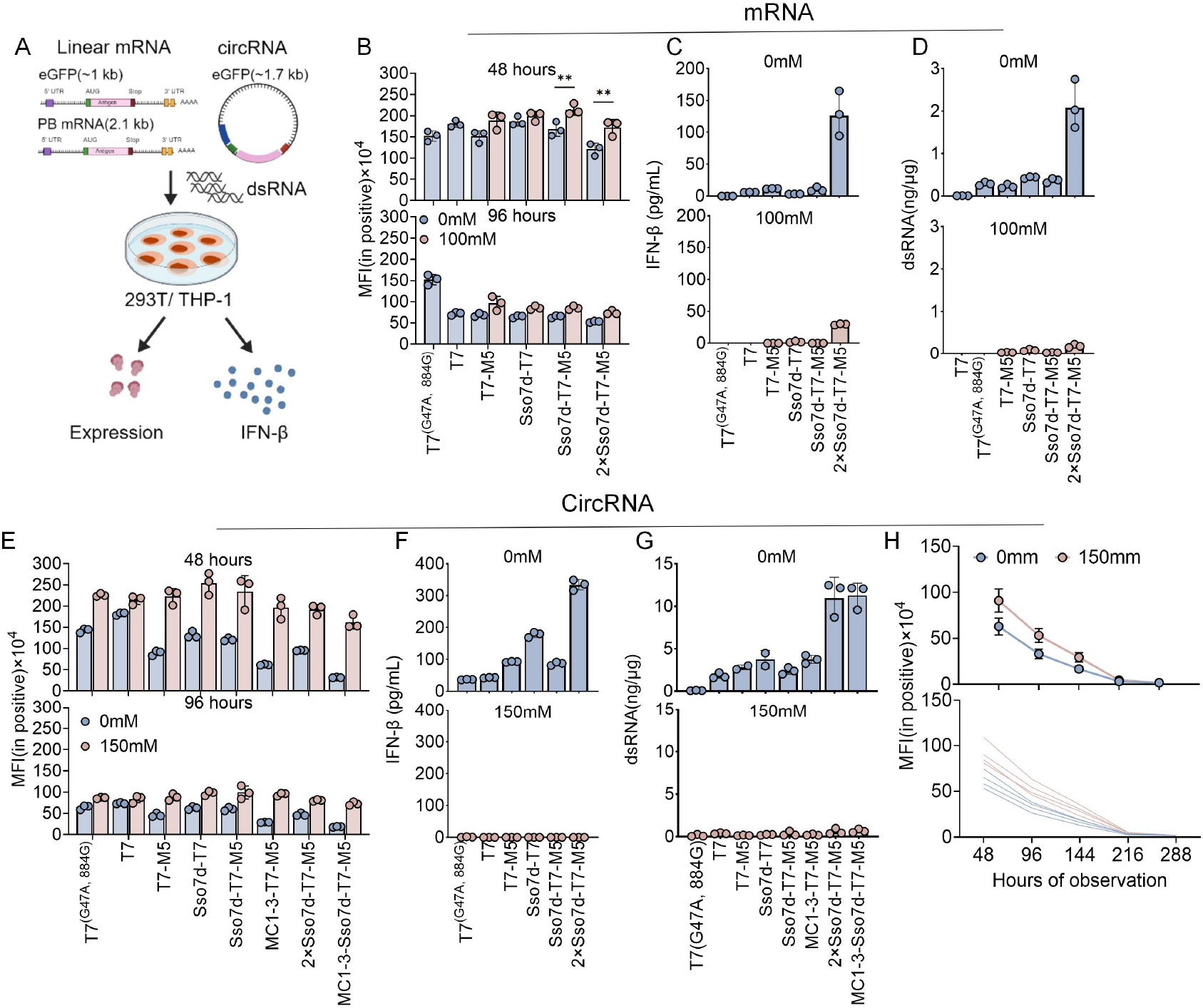
Bioactivity and immunogenicity profiling of mRNA and circRNA transcripts synthesized by engineered T7 RNAPs across a salt gradient. (A) Schematic of the experimental workflow: transcripts of varying architecture and length— including linear eGFP mRNA (∼1 kb), PB mRNA (2.1 kb), and circular eGFP RNA (circRNA, ∼1.7 kb) - were synthesized utilizing the specified RNAP mutants at 0, 100, or 150 mM NaCl. These preparations were transfected into HEK293T and THP-1 cell lines to evaluate translational efficiency and innate immune activation (via IFN-β secretion). (B, E) Mean fluorescence intensity (MFI) of eGFP expression in HEK293T cells at 48 and 96 h post-transfection for linear mRNA (B) and circRNA (E). (C, F) IFN-β secretory profiles of THP-1 cells following stimulation with linear mRNA (C) and circRNA (F) preparations. (D, G) Quantitative assessment of residual dsRNA content within linear mRNA (D) and circRNA (G) cohorts. (H) Longitudinal eGFP expression over 288 h; upper panel summarizes four circRNA mutants with distinct splicing introns, lower panels show individual trajectories. Notably, DBD-tethered chimeras maintained robust translational potency and negligible immunogenic signatures across all ionic strengths. Conversely, wild-type (WT) T7 and the T7(G47A/P884G) mutant exhibited a marked attenuation in transcriptional performance and product bioactivity under elevated salt conditions. Data are expressed as mean ±s.d. (n=3).

### Halotolerant T7 RNAP Chimeras improve circRNA Translational Efficiency and Minimize InnateImmune Activation

To delineate the causal relationship between dsRNA contaminants and innate immune activation, an IVT platform utilizing unmodified nucleotides was employed to synthesize eGFP circRNA^45^. The purity of circRNA remained robustly unaffected by high-ionic-strength conditions(Supplementary File 1: Figure S8). RNA unmodified nucleotides were specifically chosen to circumvent the immunosuppressive masking effects of pseudouridine, thereby sensitizing the system to PRR-mediated signaling triggered by dsRNA impurities. Quantitative analysis in 293T cells revealed that eGFP circRNA synthesized under high-salt conditions across all chimeric T7 RNAP mutants demonstrated significant increases in translational potency, with eGFP expression levels rising by 20–400% relative to low-salt controls (Figure 6E). Notably, translational performance converged among the distinct engineered mutants within the high-salt cohort, suggesting that a robust and generalizable correlation between high-salt transcriptional suppression of dsRNA formation and enhanced RNA expression. In contrast, circRNA generated under standard low-salt conditions induced potent IFN-β secretion (Figure 6G). A strong positive correlation was established between residual dsRNA mass fractions and IFN-β induction levels (Figure 6F). Specifically, chimeras featuring higher DBD affinities (such as homotypic or heterotypic DBD-chimeric enzymes) exhibited elevated dsRNA byproduct formation and correspondingly heightened proinflammatory signaling. These data confirm that dsRNA is the primary ligand driving innate immune activation in these systems. Conversely, circRNA synthesized via high-salt IVT, which maintained dsRNA levels below the limit of detection, failed to elicit any measurable IFN-β response (Figs. 6F, 6G). These findings indicate that the ionic strength-dependent attenuation of antisense transcription effectively abrogates innate immune recognition.

To further validate the generalizability of this platform, four additional eGFP circRNA mutants featuring distinct splicing introns were evaluated^46,47^(Plasmid sequences are provided in Supplementary File 2: Table S2). All mutants consistently demonstrated enhanced expression under high-salt conditions, achieving an average increase of 50% compared to low-salt groups (Figure 6H). These results indicate that high-ionic strength transcription not only preserves the molecular integrity of linear mRNA but exerts a disproportionately beneficial effect on the synthesis and translational performance of circRNA, particularly in the absence of modified nucleosides. Collectively, these findings suggest that the suppression of dsRNA through high-salt process optimization is an essential strategy for the high-yield production of low-immunogenicity circRNA.

### Non-Fed-Batch Process Intensification Using Halotolerant T7 RNAP Chimeras for High-Titer RNA Production

Substrate inhibition and suboptimal ionic strength represent critical bottlenecks that limit the volumetric productivity of IVT systems. Traditional bioprocessing relies on fed-batch strategies to maintain low instantaneous rNTP concentrations (typically <10 mM for standard reactions and <5 mM for fed-batch systems) to circumvent enzymatic inhibition mediated by high substrate molarity and the associated sodium counterion load^48–50^. We hypothesized that engineering T7 RNAP for enhanced halotolerance would permit elevated initial rNTP concentrations in single-batch (non-fed-batch) configurations, thereby intensifying the manufacturing process and maximizing yield.

In the PB mRNA transcription system (cumulative [Na+] = 270 mM), increasing initial rNTP concentrations from 10 mM to 15 mM resulted in significant yield attenuation across all chimeric mutants (Figure 7A). This indicates that halotolerance alone is insufficient to override the complex inhibitory effects associated with the supraphysiological substrate concentrations required for long-chain linear mRNA synthesis. In stark contrast, the Sso7d-T7-M5 chimera demonstrated exceptional substrate resilience in the circRNA synthesis system (Figure 7B). Utilizing 15 mM of each rNTP, we achieved a volumetric productivity of 15 mg/mL, representing a 50% increase over the 10 mM baseline (Figure 7B). However, further titration to 17.5 mM rNTP led to diminishing marginal returns and a subsequent decline in yield, likely due to substrate-induced catalytic stalling or osmotic stress on the elongation complex. Temporal profiling of RNA accumulation revealed that this enhancement is duration-dependent; while no significant divergence was observed during the initial 2-hour phase, the cumulative advantages of high-substrate loading became pronounced after 4 hours of reaction (Supplementary File 1: Figure S9). Furthermore, although the tandem fusion 2×Sso7d-T7-M5 tolerated 15 mM rNTP, it failed to yield synergistic productivity gains, suggesting that steric constraints or over-stabilization of the DNA-binding domain may limit turnover rates at extreme substrate concentrations. These findings demonstrate that rational engineering of RNA polymerase salt tolerance effectively mitigates rNTP substrate inhibition in unmodified nucleotide IVT systems (such as circRNA), enabling high-titer RNA synthesis (15 mg/mL) in single-batch reactions. This approach provides an enzymatic solution for streamlining manufacturing workflows and eliminating fed-batch requirements

**Figure 7.**
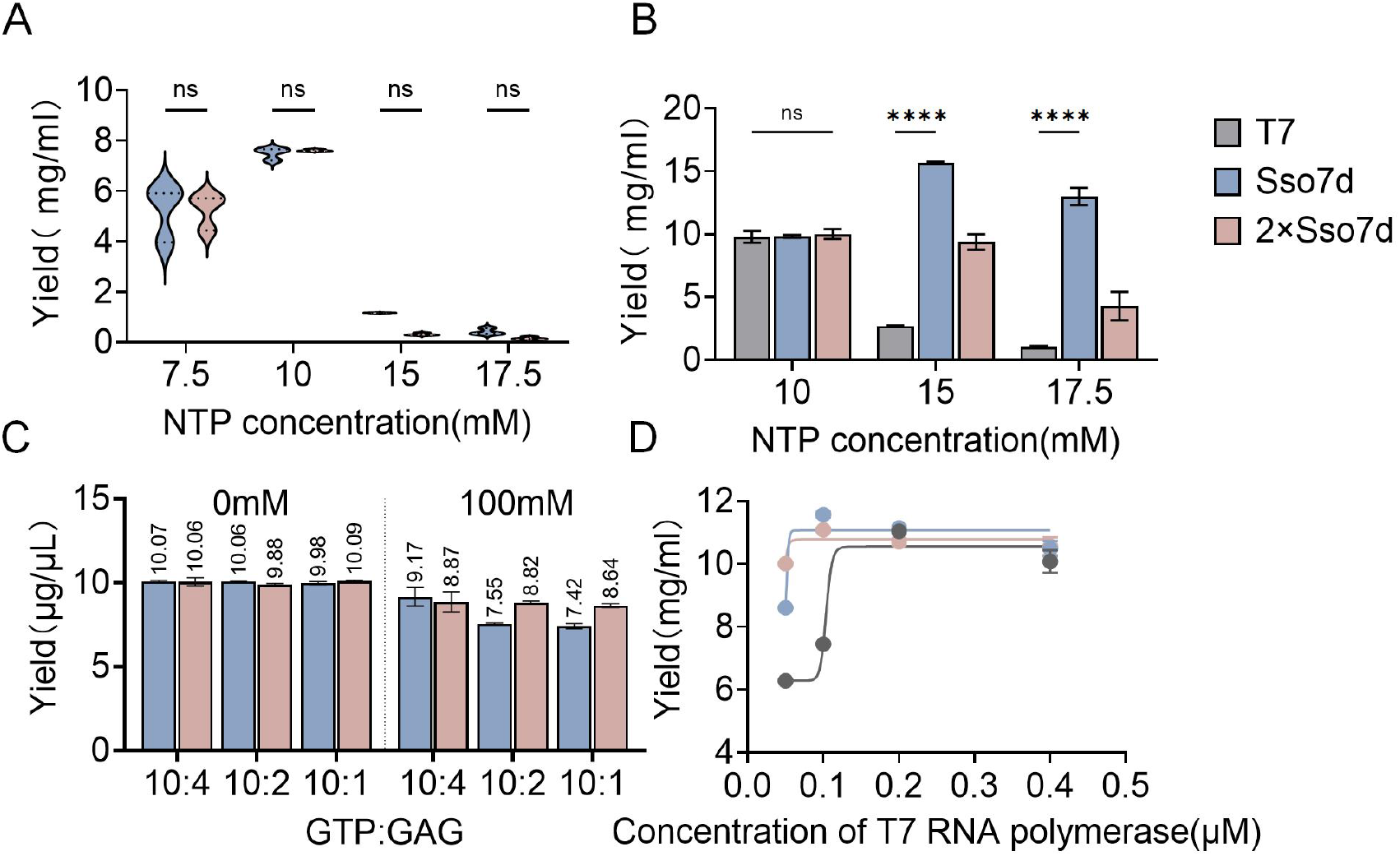
Systematic optimization of reaction parameters to maximize the specific productivity of Halotolerant T7 RNAP mutants. (A) NTP titration profiles revealing a precipitous decline in RNA yield at total nucleotide concentrations exceeding 10 mM under conventional reaction conditions. (B) Comparative analysis of substrate tolerance under high-salt. Notably, DBD-tethered chimeras sustain robust transcriptional output at hyper-concentrated NTP levels (15-17.5 mM), a regime where wild-type T7 displays near-complete catalytic inhibition. (C) Strategic optimization of the GTP:cap analog (GAG) molar stoichiometry at 0 and 100 mM NaCl for the Sso7d- and 2×Sso7d-T7-M5 iterations. (D) Enzyme dose-response curves (0-0.5 μM) demonstrating the saturation kinetics of engineered polymerases and defining the optimal enzyme-to-template ratio for maximized yield. Data are expressed as mean ± s.d. (n = 3). Statistical significance was determined via one-way ANOVA (****P < 0.0001; ns, not significant).

We further evaluated the relationship between cap analog utilization and transcriptional productivity was subsequently evaluated for the Sso7d-T7-M5 and 2×Sso7d-T7-M5 chimeras. Stoichiometric titration of cap analogs demonstrated that volumetric yield was largely independent of analog concentration under low-salt conditions for both mutants. Conversely, under high-salt conditions—specifically at GTP:GAG ratios of 10:2 and 10:1—the tandem 2×Sso7d-T7-M5 mutant exhibited superior productivity compared to the monomeric Sso7d-T7-M5 chimera, although both mutants displayed a net reduction in yield relative to their respective low-salt baselines (Figure 7C).

To elucidate the correlation between specific catalytic activity and manufacturing efficiency for long-chain mRNA, PB mRNA yields were assessed across a gradient of enzyme concentrations. The engineered chimeras, Sso7d-T7-M5 and 2×Sso7d-T7-M5, reached yield parity with wild-type T7 RNAP while requiring only 50% of the standard enzyme dosage (Figure 7D). This 2-fold increase in catalytic efficiency underscores the enhanced processivity and template affinity conferred by the DBD fusions.

Collectively, these engineered mutants facilitate robust process intensification, permitting a 50% reduction in enzyme consumption without compromising cumulative RNA output. While high-salt conditions marginally attenuate co-transcriptional capping efficiency, the synergistic gains in volumetric productivity and the near-complete suppression of immunogenic dsRNA byproducts establish a superior equilibrium between product quality, cost-effectiveness, and industrial scalability. These advancements effectively streamline the manufacturing workflow by mitigating the need for complex fed-batch operations.

## Discussion

IVT has emerged as a cornerstone of RNA-based biomanufacturing. Kinetic frameworks, supported by experimental validation, categorize IVT regimes as either initiation-limited or elongation-limited, dictated by the prevailing solution conditions^51^. High-salt (e.g., NaCl, Mg^2+^) typically attenuates transcription by destabilizing the initial binding affinity between the polymerase and the promoter. In the context of the “scrunching” model—where T7 RNAP remains anchored to the promoter while pulling in downstream DNA—the transition from unstable initiation complexes to stable elongation complexes represents a high-energy barrier. High salt concentrations exacerbate this instability, often leading to abortive cycling and reduced promoter escape efficiency.

This study demonstrates that the synergy between catalytic core stabilization (T7-M5) and optimized N-terminal DBD fusions effectively lowers this barrier. While the M5 mutations preserve the global structural integrity of the polymerase under ionic stress, the heterologous DBDs function as molecular anchors. By enhancing promoter recruitment and stabilizing the transcription bubble during the stressed scrunching phase, these chimeras facilitate a more robust transition to the EC even under high salt conditions. This engineering strategy not only preserves volumetric productivity but also significantly suppresses the formation of immunostimulatory dsRNA. This effect is particularly pronounced in circRNA production using unmodified nucleotides, where the suppression of antisense transcription is paramount for therapeutic safety. Mechanistically, our data suggest that chimeric mutants leverage high-salt-induced electrostatic shielding to regulate the enzyme’s processivity. During elongation, high-salt weakens non-specific RNA–polymerase interactions, thereby inhibiting the RdRP activity responsible for dsRNA synthesis. Furthermore, the precise calibration of DBD-mediated affinity ensures that terminal release efficiency is maintained. Unlike conventional high-salt systems that suffer from severe productivity losses, our platform achieves a kinetic equilibrium: it provides sufficient “grip” for initiation and processive elongation while allowing for the precise dissociation required for high 3’ homogeneity.

We acknowledge that certain heterotypic tandem chimeras, despite their high specific activity, retain an excessive DNA-binding affinity that is not entirely alleviated by high-salt. Such residual “over-binding” accounts for marginally higher dsRNA levels compared to the lead Sso7d-T7-M5 mutant. Although Sso7d-T7-M5 exhibited slightly lower 3’-homogeneity than the tandem mutants, its superior ability to ablate dsRNA contaminants to below 0.001% meets the most stringent industrial requirements. Future iterations focus on the calibration of DBD binding affinity or the integration of synergistic mutations to attenuate dsRNA formation to further refine these chimeric scaffolds. In conclusion, this dual protein-and-process engineering strategy provides a scalable, cost-effective solution for high-titer, high-purity RNA manufacturing. By circumventing the need for complex fed-batch operations and arduous downstream dsRNA removal, this platform streamlines the production of both research-grade and clinical-grade RNA therapeutics, fundamentally advancing the feasibility of synthetic biology applications.

## Materials and Methods

### Cell Culture

CHO (Chinese hamster ovarian carcinoma) cells were obtained from ATCC and grown in Ham’s F-12K (Kaighn’s) Medium (F12K; Gibco, USA) supplemented with 10% FBS (Gibco, USA), 1% L-Glutamine (Gibco, USA), 1% HT Supplement (Gibco, USA) and 1% Penicillin-Streptomycin (P/S; ThemoFisher, USA). HEK293T (Human Embryonic Kidney 293T cells) cells were obtained from ATCC and grown in Dulbecco’s Modified Eagle Medium (DMEM; Corning, USA) supplemented with 10% FBS (Gibco, USA) and 1% Penicillin-Streptomycin (P/S; ThemoFisher, USA). THP-1 cells were obtained from ATCC and grown in Modified Eagle Medium (RPMI-1640; ATCC, USA) supplemented with 10% FBS (Gibco, USA), 0.05 mM β-mercaptoethanol and 1% Penicillin-Streptomycin (P/S; ThemoFisher, USA). All cells were maintained at 37°C in a humidified atmosphere containing 5% CO2.

### Plasmid Construction

The plasmids encoding T7 RNAP and its mutants were commercially synthesized by General Biosystems (Chuzhou, China). Briefly, synthetic genes encoding the polymerase and its mutated derivatives (harboring substitutions such as P266L, S430P, S643P, F849I, F880Y, and Q744R) were codon-optimized and ligated into the pQE80L vector (Qiagen, Germany). This expression system, under the control of a T5 promoter, was configured to incorporate an N-terminal hexahistidine (6×His) tag to facilitate subsequent affinity purification. Following the identification of lead mutant candidates, DNA sequences encoding diverse DNA-binding domains (DBDs) from the MC1, Sul7d, and Cren7 families were synthesized and fused to the NTD of the T7 RNAP. To modulate the spatial orientation of these heterologous domains and maintain the structural integrity of the T7 RNAP, flexible linkers with the amino acid sequence (GGGGS)_3_ were orchestrated both between the 6×His tag and the DBD, and between the DBD and the polymerase. The resulting chimeric constructs represent a comprehensive repertoire of engineered polymerases with N-terminal fusions. Complete plasmid sequences are provided in Supplementary File 2: Table S1.

Concurrently, DNA templates for the IVT of PiggyBac (PB) mRNA, eGFP mRNA and eGFP circular RNA (circRNA)—all driven by a T7 promoter—were synthesized by General Biosystems. Detailed sequence information for these transcription templates is archived in Supplementary File 2: Table S2.

### Protein expression and purification

#### Expression of T7 RNA Polymerase

The recombinant plasmids encoding T7 RNAP mutants were transformed into Escherichia coli BL21(DE3) competent cells (Weidi Biotechnology, China). The transformants were cultured overnight at 37°C in LB medium supplemented with 100 μg/mL ampicillin. Upon reaching an optical density at 600 nm (OD_600_) of 0.6–0.8, protein expression was induced by the addition of isopropyl β-D-1-thiogalactopyranoside (IPTG) to a final concentration of 1 mM. Following a 4-hour induction period at 37°C, the cells were harvested via centrifugation.

#### Purification of T7 RNA polymerase

The cell pellets were resuspended in lysis buffer (50 mM Tris-HCl, pH 7.5, 500 mM NaCl, 5 mM imidazole, 1 mM PMSF) and disrupted by ultrasonication. The resulting lysate was clarified by centrifugation, and the supernatant was subsequently passed through a 0.22 μm filter. Initial capture was performed using Ni-NTA affinity chromatography (Purose 6 FF, Qianchun Biology, China). The column was equilibrated with binding buffer, and the target proteins were eluted using a linear imidazole gradient (100– 300 mM). Fractions displaying >70% purity were pooled, and their ionic strength was adjusted to facilitate subsequent polishing. Fine purification was achieved through SP cation-exchange chromatography (Purose 6 HP, Qianchun Biology). The column was equilibrated with 50 mM Tris-HCl (pH 7.5) and 100 mM NaCl. After loading the captured fractions, elution was orchestrated using a linear salt gradient ranging from 100 mM to 500 mM NaCl in 50 mM Tris-HCl (pH 7.5). Fractions with >85% purity were collected, concentrated via ultrafiltration, and dialyzed overnight against a storage buffer (25 mM Tris-HCl, pH 7.5, 100 mM NaCl, 25 mM β-ME, 0.1% Triton X-100, 50% glycerol).

### Measurement of RNA Polymerase Specific Activity

iSpinach-D5 transcription templates were generated by PCR amplification of the plasmid using M13F (5’-gtaaaacgacggccagt-3’) and M13R (5’-caggaaacagctatgac-3’) universal primers. Real-time transcriptional activity of chimeric and mutant T7 RNA polymerases was monitored based on iSpinach aptamer-DFHBI binding-induced fluorescence^31,40–43^.

The standard IVT reaction mixture comprised 0.04 μg/μL RNA polymerase, 40 mM Tris-HCl (pH 8.0), 50 mM MgCl_2_, 10 mM DTT, 2 mM spermidine, 10 mM rNTPs, 16 nmol/mL DFHBI, 20 pmol/mL iSpinach-D5 DNA template, and 1 U/μL RNase inhibitor. To evaluate the salt resilience of the engineered polymerases, two distinct ionic strength conditions were tested: a standard buffer and a high-salt buffer supplemented with an additional 100 mM NaCl. Fluorescence intensity, reflecting the accumulation of functional RNA aptamers during the elongation phase, was recorded in real-time using an EnVision (PerkinElmer, USA) microplate reader at excitation/emission wavelengths of either 490/516 nm or 472/507 nm in 96-well formats.

### In vitro transcription of mRNA

Standard IVT reactions were was orchestrated in a buffer comprising 40 mM Tris-HCl (pH 8.0), 10 mM DTT, and 2 mM spermidine (Sigma-Aldrich, USA). The nucleotide pool consisted of 10 mM of each rNTP (Hongene, China); for the synthesis of modified transcripts, UTP was entirely substituted with pseudouridine triphosphate (pseudo-UTP, Hongene). The reaction mixture was further supplemented with 20 mM MgCl_2_, 0.06 mg/mL of DNA template linearized via Nhe I (BestEnzyme, Lianyungang, China), 0.04 mg/mL (0.4 µM) of T7 RNA polymerase or its engineered mutants, 0.01 μL/μL (200 U/mL) RNase inhibitor, and 0.6 U/mL inorganic pyrophosphatase (Novoprotein, China). While the basal Na^+^ concentration ranged from 120 to 160 mM, high-salt conditions were achieved by supplementing the system with an additional 100–200 mM NaCl. For reactions utilizing the T7 RNAP (G47A, 884G) mutant, rNTP concentrations were reduced to 4 mM each to optimize processivity and yield.

#### Enzymatic capping transcription (for PiggyBac transposase mRNA)

Following IVT incubation at 37°C for 2-4 hours, the DNA template was removed by digestion with halotolerant DNase I-ST (BestEnzyme, China). Uncapped RNA was recovered through LiCl precipitation and subsequently subjected to enzymatic capping. The capping assembly contained 0.5 mg/mL uncapped RNA, 1× Capping Buffer, 1 mM GTP, 0.2 mM S-adenosylmethionine (SAM), RNase inhibitor (Vazyme, Nanjing, China), 1 mM Vaccinia Capping Enzyme (VCE, Vazyme), and 5 mM 2’-O-methyltransferase (2’-O-Mtase, Vazyme). After a 1-hour incubation at 37°C, the capped mRNA was purified using the Monarch RNA Purification Kit (NEB, USA).

#### Co-transcriptional capping (for eGFP mRNA)

For eGFP mRNA, co-transcriptional capping was facilitated by the inclusion of the CAP GAG analog [(3’OMe) m7(3’OMeG)(5’)ppp(5’)(2’OMeA)pG] (Syngenebio, Nanjing, China). The polymerase concentration was increased to 0.075 mg/mL (0.75 µM). Other reaction parameters remained consistent with the standard IVT protocol.

#### Circular RNA IVT (eGFP circRNA)

Reactions were performed using the standard system with modifications, including 40 mM MgCl_2_, 0.1 mg/mL DNA template, and the exclusive use of unmodified natural rNTPs. Post-transcriptional enrichment of circRNA was achieved via RNase R digestion (GenScript, China), followed by purification using Monarch RNA purification kits (NEB, USA). All RNA products were quantified utilizing a NanoDrop™ One spectrophotometer (Thermo Scientific™, USA).

### RNase T1 digest assay

#### Preparation of Transcription Templates

Sense and antisense long single-stranded DNA oligonucleotides corresponding to the transcription templates described by Dousis et al.^20^ (Supplementary File 2: Table S3), were commercially synthesized by Genewiz (Suzhou, China). These oligonucleotides were annealed at a 1:1 molar ratio in annealing buffer by heating at 95°C for 5 min, followed by gradual cooling to 25°C over 30 min. The successful formation of double-stranded DNA (dsDNA) was validated via agarose gel electrophoresis. RNA samples were subsequently synthesized using the baseline in vitro transcription (IVT) system.

#### RNase T1 Mapping and LC-MS Analysis

RNase T1 digestion was performed according to the methodology established by Jiang et al.^44^, leveraging the enzyme’s capacity to selectively cleave 3’ phosphodiester bonds at guanosine residues within single-stranded RNA regions. Briefly, mRNA (1 mg/mL, 40 μL) was mixed with 8 M urea (60 μL; Sigma-Aldrich, Germany), 1 M Tris-HCl at pH 7.0 (12 μL; Invitrogen, USA), and 0.5 M EDTA (0.8 μL; Invitrogen, USA). Following denaturation at 90°C for 10 min and subsequent cooling to room temperature, the mixture was treated with RNase T1 (1000 U/μL, 20 μL; Thermo Fisher Scientific, USA) and incubated at 37°C for 15 min. The resulting digestion fragments were resolved via reversed-phase ion-pair high-performance liquid chromatography (RP-IP-HPLC) using an ACQUITY Premier Oligonucleotide BEH C18 column (2.1 × 100 mm, 1.7 μm). Qualitative and quantitative characterizations were conducted on an LC-MS system integrated with a Waters Xevo G2-XS QTof high-resolution mass spectrometer (Waters, USA).

### Purity assessment of transcription products

RNA integrity was assessed using the Agilent Fragment Analyzer 5300 system. Briefly, RNA samples were diluted to a final concentration of 0.1 mg/mL and analyzed with the Agilent DNF-471 RNA Kit (15 nt) (Agilent Technologies, USA) in accordance with the manufacturer’s protocols. Full-length product (FLP) profiles were subsequently generated and quantified using ProSize Data Analysis Software (Agilent Technologies, USA) to evaluate the proportion of intact transcripts and the extent of premature termination.

### Capping efficiency analysis

To characterize the 5’-terminal capping efficiency of the target mRNA (5’UTR: AGGGAAGAAGAAAGAAAGGAAAGUAAACCACCCACGGACCUCA), a 5’-biotinylated DNA probe (5’-CY5-dTdGdAdGdGdTmCmCmGmUmGmGmGmUmGmGmUmUmUmAmCmUmUmUmCmCm U-3’ Biotin) was strategically designed. The mRNA and probe were mixed at a 1:1 molar ratio, denatured at 85°C for 1 min, and subjected to a step-wise gradient annealing (65°C, 55°C, and 40°C for 2 min each) until reaching 22°C to facilitate the formation of a stable RNA/DNA hybrid. The annealed complex was subsequently incubated with thermostable RNase H (5 U; NEB, USA) at 37°C for 2 h to specifically hydrolyze the RNA strand within the hybrid region, thereby releasing 5’-oligonucleotide fragments containing diverse capping states (e.g., Cap0, Cap1, or G-cap). These 5’ fragments were captured using streptavidin-coupled magnetic beads (Dynabeads MyOne Streptavidin C1, 10 mg/mL; Invitrogen, USA) via a 30-min incubation at room temperature. Following stringent washing steps (twice with 2× B&W buffer and twice with nuclease-free water), the captured fragments were robustly eluted in 1 mmol/L EDTA containing 1% methanol by heating at 85°C for 3 min. The supernatant was recovered after centrifugation at 12,000 rpm for 10 min. Analytical characterization was performed using a Waters Ultra-Performance Liquid Chromatography-Mass Spectrometry (UPLC-MS) system. The fragments were resolved on an ACQUITY Premier Oligonucleotide BEH C18 column (2.1 × 100 mm, 1.7 μm) maintained at 65°C with a flow rate of 0.3 mL/min. The mobile phase system consisted of an aqueous solution of 0.75% HFIP, 0.0375% DIEA, and 10 μM EDTA (A) and acetonitrile (B), employing a linear gradient (0–10 min, 5%–50% B; 10.5–12.5 min, 90% B). Mass spectrometry was operated in negative ion full-scan mode (m/z 400–3000) with a capillary voltage of 2.5 kV and a desolvation temperature of 400°C. The capping efficiency was determined by integrating the peak areas from the extracted ion chromatograms (EICs) of various terminal forms (uncapped ppp-, pp-, p-, Cap0, and Cap1) via the area normalization method.

Enzymatic capping transcription: Capping efficiency 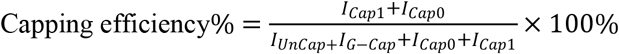

Co-transcriptional capping: Capping efficiency 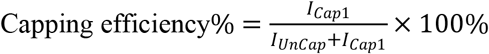

*I* denotes peak intensity

### dsRNA ELISA assay

The concentration of residual double-stranded RNA (dsRNA) contaminants in IVT mRNA was determined using a sandwich ELISA. A commercial dsRNA quantification kit (Cat. No. RES-A092, ACRO Biosystems, China) was employed, utilizing a 400-bp unmodified dsRNA as the reference standard. A standard curve, encompassing a concentration range of 25–800 pg/mL, was established via serial dilution. Both mRNA samples and standards (100 μL per well) were added to 96-well plates pre-coated with the anti-dsRNA capture antibody (K1) and incubated for 60 min at room temperature (RT, 25°C) with agitation at 400 rpm. Following four successive washes with wash buffer, horseradish peroxidase (HRP)-conjugated anti-dsRNA detection antibody (J2, 0.5 μg/mL) was introduced and incubated under identical conditions. After a final four-wash cycle, TMB substrate was added and allowed to develop for 20 min in the dark to facilitate chromogenic reaction before being quenched with stop solution. Absorbance values at 450 nm and 630 nm were recorded using a Spark multimode microplate reader (Tecan, Switzerland). Sample concentrations were modulated and calculated by fitting the standard curve with a four-parameter logistic (4-PL) regression model. All assays were performed in duplicate, utilizing dilution buffer as a negative control; a coefficient of determination (R^2^ ≥ 0.99 was underscored as the prerequisite for a valid standard curve.

### In vitro activity assay

#### eGFP reporter assay

HEK293T cells were maintained for 36 h and subsequently seeded into 48-well plates at a density of 5×10^5^ cells per well. Upon reaching 90% confluency, the cells were transfected with 1 μg of capped mRNA utilizing Lipofectamine 3000 (Vazyme, China) in accordance with the manufacturer’s instructions. Briefly, 1 μg of mRNA was diluted in 50 μL Opti-MEM and supplemented with 2 μL of P3000 reagent; this mixture was then combined with an equal volume of diluted Lipofectamine 3000. Following a 15 min incubation at room temperature, the resulting lipoplexes were added to the cell culture. After 24 h of incubation at 37°C, the cells were washed with PBS, dissociated with 0.5% Trypsin-EDTA, and analyzed via flow cytometry to determine both the mean fluorescence intensity (MFI) and the percentage of EGFP-positive cells.

#### PiggyBac (PB) transposase activity assay

A reporter plasmid (Ai9-PB) was constructed based on the Ai9 architecture originally described by Madisen et al.^52^. This construct features a CAG promoter-driven tdTomato expression cassette interrupted by a 3×SV40 polyA stop signal. To facilitate PB-mediated transposition, the LoxP elements in the original Ai9 vector were replaced with PiggyBac-specific inverted terminal repeats (ITRs) (sequences provided in Supplementary File 2: Table S4. This reporter system was integrated into the CHO cell genome via electroporation to establish a stable reporter cell line. For the transposase activity assay, 0.2 μg of PB mRNA was transfected into 4×10^5^ reporter cells using the Lonza Nucleofector system (Program EH-133; Lonza, Switzerland). On day 6 post-transfection, cells were harvested and analyzed via flow cytometry. The percentage of tdTomato-positive cells and the MFI were recorded to quantitatively evaluate the translation efficiency and functional potency of mRNA transcripts synthesized by the diverse T7 RNAP mutants.

### In vitro immunogenicity assessment

HEK293T or THP-1 cells were seeded at 5×10^5^ cells per well in 48-well plates (NEST, China) and incubated at 37°C under 5% CO_2_ for 24 h to achieve 70-80% confluence. Prior to transfection, culture medium was exchanged for serum-free Opti-MEM. Test mRNA (1μg) and Lipofectamine 3000 (2 μL; Vazyme, China) were separately diluted in 50 μL Opti-MEM, incubated at room temperature for 5 min, mixed at a 1:1 ratio, and further incubated at room temperature for 15 min to permit liposome–RNA complex formation. The transfection mixtures were added to the cells, and medium was replaced with complete culture medium after 6 h.

Experimental groups included a positive control (142-bp dsRNA, 50 ng per well; Jena Bioscience, Germany), a negative control (transfection reagent alone), and a blank control. At 24 h post-transfection, culture supernatants were harvested and clarified by centrifugation at 300× g for 5 min at 4 °C to pellet cellular debris. IFN-β levels in the supernatants were quantified using a human IFN-β ELISA kit (Abclone, China) according to the manufacturer’s instructions. All experiments were performed with three technical replicates per group and independently repeated at least three times.

### Statistical analysis

Statistical analyses were carried out using GraphPad Prism 10.0 (GraphPad Software, San Diego, CA). Data were analyzed using unpaired *t* tests to compare two different conditions and analysis of variance for more conditions. All error bars show the standard errors of the means (SEM). Statistics significant difference was considered as following: P≥ 0.05 (ns), P < 0.05 (*), P < 0.01 (**), P < 0.001 (***), P<0.0001 (****).

## Acknowledgements

This study was supported by Shanghai Science and Technology Innovation Action Plan, Cell and Gene Therapy Project, Science and Technology Commission of Shanghai Municipality (23J21900500).

